# Emotion Dysregulation Links Interoceptive Accuracy and Vagal Activity to Psychopathology in Male but not Female Adolescents

**DOI:** 10.64898/2026.09.14.751467

**Authors:** Maximilian Schmausser, Hanna Schwinde, Ines Mürner-Lavanchy, Corinna Reichl, Michael Kaess, Julian Koenig

## Abstract

**Background:** Emotion dysregulation is common across psychiatric disorders, but the biological mechanisms underlying emotional vulnerability remain poorly understood. Cardiac vagal regulation and interoception, the perception of bodily signals, may jointly contribute to emotion regulation. We examined whether their interaction is associated with emotion dysregulation and psychopathology in adolescents receiving psychiatric inpatient treatment.

**Methods:** We studied 279 adolescent psychiatric inpatients. Cardiac vagal regulation was assessed using heart rate variability and interoceptive accuracy using a measure of bodily signal perception. Structural equation modeling tested associations between cardiac vagal regulation, interoceptive accuracy, emotion dysregulation, depressive symptoms, and personality dysfunction, including sex-specific pathways and interactions.

**Results:** Here we show that associations between autonomic regulation, interoception, and psychopathology differ by sex. In male adolescents, lower cardiac vagal regulation is associated with greater emotion dysregulation, which is associated with higher depressive symptoms and greater personality dysfunction. These associations depend on interoceptive accuracy and are most evident at low to intermediate levels of bodily signal perception. Comparable pathways are absent in female adolescents.

**Conclusions:** These findings indicate that interactions between cardiac regulation and bodily signal perception contribute to emotional vulnerability in a sex-dependent manner. Altered body–brain communication may contribute to divergent developmental pathways of psychopathology. Integrating sex-specific autonomic and interoceptive processes may improve models of emotion regulation and psychiatric risk during adolescence.

**Plain Language Summary:** Difficulties in regulating emotions are common across mental health conditions, but the biological factors underlying vulnerability remain poorly understood. This study investigated whether heart activity and the ability to perceive bodily signals are linked to emotional and mental health problems during adolescence. We examined 279 adolescents receiving psychiatric inpatient treatment and assessed their heart activity, perception of bodily signals, emotion regulation, and psychological symptoms. In male adolescents, lower vagal regulation of the heart was linked to greater emotion dysregulation, which was associated with more depressive symptoms and personality dysfunction. These relationships were strongest among those with lower to moderate accuracy in perceiving bodily signals and were not observed in females. The findings suggest that heart–brain communication may contribute to emotional and psychological vulnerability during adolescence, with potential implications for sex-specific prevention and treatment.

## 1. Introduction

Mental disorders constitute a major public health burden across the lifespan and are among the leading causes of disability worldwide. Across both internalizing and externalizing forms of psychopathology, disturbances in emotional functioning represent a central clinical feature.^1^ To better characterize the broad range of emotional difficulties observed across psychiatric disorders, the construct of *emotion dysregulation* has increasingly emerged as a transdiagnostic framework spanning childhood, adolescence, and adulthood. Emotion dysregulation has commonly been defined as a pattern of emotional experience, expression, and regulation that interferes with adaptive, goal-directed behavior.^2^ Importantly, emotion dysregulation does not reflect a unitary deficit localized to a single process but rather can emerge across multiple stages of emotional functioning and regulation. These stages include the identification of emotional states, the selection of regulatory strategies, the implementation of regulatory actions, and the ongoing monitoring and adjustment of regulatory processes. Across these stages, impairments may arise at the level of perception, valuation, or action, thereby contributing to maladaptive emotional responding in distinct ways.^3^ Consequently, emotion dysregulation encompasses a broad range of impairments, including heightened emotional reactivity, difficulties in identifying and differentiating emotions, reduced emotional awareness and acceptance, maladaptive evaluations of emotional states, impaired selection or implementation of adaptive regulation strategies, deficits in inhibitory control under emotional distress, and difficulties flexibly adapting emotional responses to changing contextual demands.^4^ A growing body of evidence supports the central role of emotion dysregulation across a wide range of psychiatric disorders, including depression, anxiety disorders, borderline personality disorder, eating disorders, substance use disorders, and neurodevelopmental disorders.^5,6^ Difficulties in emotion regulation have consistently been associated with greater symptom severity, functional impairment, self-injurious behavior, and poorer treatment outcomes.^7,8^ Particularly during adolescence, which represents a developmental period characterized by profound neurobiological, hormonal, autonomic, and socio-emotional changes, deficits in emotion regulation appear to constitute a major vulnerability factor for the emergence and persistence of affective psychopathology.^2^ Despite its clinical importance, however, the neurobiological and psychophysiological mechanisms underlying individual differences in emotion dysregulation remain incompletely understood.

Contemporary models of emotion and emotion regulation increasingly emphasize the role of body–brain interactions in shaping emotional experience and adaptive regulatory functioning. Rather than viewing emotions as solely brain-based phenomena, these accounts propose that the perception, integration, and regulation of bodily states contribute fundamentally to how emotions are generated, differentiated, and regulated.^9,10^ In this framework, emotional experiences emerge through the continuous integration of physiological signals from the body with contextual and conceptual information, thereby enabling the organism to flexibly adapt behavior to changing environmental and internal demands.^11^ Consequently, individual differences in the processing and regulation of bodily signals may represent an important mechanism underlying variability in emotion regulation capacities and vulnerability to psychopathology.

The processing, sensation, perception, and awareness of visceral signals transmitted via the autonomic nervous system (ANS) to the central nervous system (CNS) is commonly referred to as *interoception.*^12^ Through hierarchically organized feedback loops, interoceptive processing continuously informs the individual about its internal state thereby contributing to the regulation of emotional, cognitive and behavioral responses.^13–15^ A growing body of evidence suggests that interoception plays a crucial role in emotion regulation and affective psychopathology. By providing a continuous stream of information about the body’s internal milieu, interoceptive processing supports the generation, differentiation, and modulation of emotional states.^16–20^ Higher interoceptive capacities have been associated with more precise emotion differentiation, ^21–23^ greater use of adaptive emotion regulation strategies, such as reappraisal ^24–26^, and stronger alignment between subjective affect and physiological responses. ^27–29^ Conversely, reduced interoceptive sensitivity has been associated with difficulties in emotional awareness and regulation. ^30,31^ In line with these findings, alterations in interoceptive processing have been identified as a transdiagnostic feature across a range of mental disorders, ^32–35^ including depression, ^10^ anxiety, ^36^ eating and substance use disorders, ^33,37^ and personality disorders,^38^ supporting the notion that interoceptive dysfunction may represent a transdiagnostic mechanism contributing to emotion dysregulation and psychopathology. Interestingly, interoception exhibits significant sex differences, with women generally showing lower interoceptive accuracy across multiple modalities compared to men, despite outperforming men on measures of emotional processing and self-awareness.^26,39^ These discrepancies suggest that the mapping between bodily signals and subjective emotional experience may differ as a function of sex, potentially contributing to sex-specific vulnerability patterns in affective psychopathology.

Among the key pathways linking the brain and body, the vagus nerve constitutes a principal component in interoceptive signaling and autonomic regulation. Through extensive afferent and efferent connections between the brain and other inner organs, the vagus nerve allows the brain not only to sense but to regulate bodily states, adjusting internal physiology in accordance with changing environmental, motivational, and emotional demands. Vagally mediated heart rate variability (vmHRV) provides a widely used, noninvasive index of (cardiac) vagal activity. Dynamic fluctuations in vmHRV accompany affective states typically decreasing during high-arousal or negative emotions and increasing during calm or positive affective states. ^40–42^ Beyond momentary emotional reactivity, resting vmHRV has been linked to emotional experience, ^43^ flexibility, ^44,45^ and regulation, with lower vmHRV associated with greater difficulties in emotion regulation as measured by the Difficulties in Emotion Regulation Scale (DERS).^46–48^ Moreover, chronically reduced vmHRV has been identified as a transdiagnostic marker of affective vulnerability, associated with an increased risk for affective disorders, ^49–51^ as well as with other disorders characterized by pronounced emotion dysregulation, such as borderline personality disorder. ^52^ Similar to interoceptive processing, vmHRV exhibits pronounced sex differences across development. During childhood and adolescence, girls typically show lower vmHRV than boys, whereas adult women generally display higher vmHRV than men, despite their elevated risk for affective disorders.^53,54^ This developmental shift suggests that autonomic and interoceptive mechanisms are dynamically modulated across the lifespan, with potential implications for sex-specific trajectories of emotion regulation and vulnerability to psychopathology.

Taken together, previous findings point to a close interrelation between cardiac vagal activity, interoceptive processing, emotion regulation capacities, and psychopathology. Yet, despite converging evidence linking interoceptive processing and vmHRV separately to emotion regulation and psychopathology, few studies have examined these processes simultaneously within a transdiagnostic developmental clinical framework. In particular, the extent to which interoceptive processing and cardiac vagal activity jointly contribute to emotion dysregulation in adolescent psychiatric populations, as well as their potential modulation by sex, remains insufficiently understood. To address this gap, we examined these relationships in a clinical sample of adolescent inpatients exhibiting varying levels of affective symptoms and emotion regulation difficulties. Drawing on prior theoretical and empirical work, we proposed that cardiac vagal activity, indexed by vmHRV, together with interoceptive processing, assessed via a heartbeat detection task, jointly contribute to emotion (dys-)regulation, measured by the DERS. In turn, emotion (dys-)regulation was expected to mediate the effects of vagal activity and interoception on psychopathological symptom severity. Given documented sex differences in interoception, vmHRV, and psychopathology, we further explored whether these associations differ between girls and boys, potentially reflecting sex-specific developmental trajectories in body–brain–emotion interactions in psychiatric inpatients.

## 2. Methods

### Participants

Data were collected within the framework of the Bernese Basis Documentation (BeBaDoc) study, which was conducted between November 2018 and December 2022 at the University Hospital for Child and Adolescent Psychiatry and Psychotherapy in Bern, Switzerland. The study (ID: 2018-01339) was approved by the Swiss Ethics Committee of the Canton of Bern and was conducted in accordance with the Declaration of Helsinki.

Study participants were adolescent inpatients aged between 11 and 18 years who were consecutively recruited within approximately four weeks (M = 27.05 days, SD = 2.07 days) following hospital admission. All participants provided written informed consent. For participants younger than 14 years, additional written informed consent was obtained from a legal guardian prior to inclusion. Exclusion criteria comprised inadequate German language skills, insufficient understanding of study details due to a clinical condition or other factors, ongoing emergency care, or lack of informed consent (by the participant and/or legal guardian). Participants received a compensation of 20 Swiss Francs for their participation.

The initial sample consisted of N = 375 participants. Of these, n = 86 participants (22.9%) were excluded due to missing data for the heartbeat detection task, n = 5 (1.3%) because of missing STiP-5.1 interview data, n = 4 (1.0%) because of missing DERS data, and one participant (0.2%) was excluded owing to an implausibly high vmHRV value (RMSSD = 314.9 ms). The final sample included N = 279 participants.

### Experimental Design

Each participant completed a single on-site session lasting approximately three hours in a designated laboratory setting. To minimize circadian influences on physiological measures, sessions were scheduled preferentially in the morning whenever feasible. The sessions began with a roughly 10-minute acclimatization period, during which participants received comprehensive information about the study procedures. Afterwards, demographic (e.g., age, sex, school type, etc.), anthropometric (i.e., height, weight, BMI) and physiological (ECG) data were collected, followed by a series of standardized psychiatric assessments. Semistructured diagnostic interviews were administered by trained clinical psychologists and psychology students, while participants independently completed self-report questionnaires. All data were recorded and managed using the web-based REDCap platform (https://www.project-redcap.org/).^87^ To ensure diagnostic reliability and adherence to quality standards, a subset of sessions was audio-recorded and subsequently reviewed by experienced clinical research personnel to assess clinical rater-agreement in diagnostics.

### Emotion dysregulation

Emotion (dys-)regulation was assessed using the Difficulties in Emotion Regulation Scale.^4^ The DERS comprises 36 items rated on a 5-point Likert scale (1 = almost never to 5 = almost always; e.g., “When I’m upset, I have difficulty controlling my behaviors”) and captures six dimensions of emotion dysregulation: (i) nonacceptance of emotional responses (*α* = 0.87), (ii) difficulties engaging in goal-directed behavior under negative affect (*α* = 0.84), (iii) difficulties controlling impulsive behavior when distressed (*α* = 0.80), (iv) lack of emotional awareness (*α* = 0.73), (v) limited access to effective emotion regulation strategies (*α* = 0.79), and (vi) lack of emotional clarity (*α* = 0.79). The DERS demonstrated excellent internal consistency in the present sample (*α* = 0.88).

### Personality pathology

The Semi-structured Interview for Personality functioning DSM-5 (STiP-5.1) is a semi-structured clinical interview designed to operationalize the DSM-5 Section III Level of Personality Functioning Scale (LPFS) as part of the Alternative Model for Personality Disorders (AMPD)^92^. The LPFS conceptualizes impairment of personality functioning across 12 facets, which are organized into four elements - identity, self-direction, empathy, and intimacy - each mapping onto one of the two overarching domains of self-functioning and interpersonal functioning. The LPFS distinguishes five levels of impairment severity (none, mild, moderate, severe, and extreme). A STiP-5.1 mean score of ≥2 (moderate impairment) is typically used as a threshold indicative of the presence of a personality disorder (PD).^92–94^ As the manual does not prescribe a specific algorithm for determining the threshold, it remains at the clinician’s discretion. In the present study, a PD was considered present when at least two of the four LPFS elements showed moderate impairment (≥2), with an overall mean of ≥2 across the four elements. The STiP-5.1 has demonstrated strong validity, reliability, and clinical utility as a dimensional measure of personality functioning.^92,95^

### Depression

Depressive symptoms were assessed using the Children’s Depression Rating Scale-Revised (CDRS-R-R),^96^ a clinician-administered interview designed to evaluate the severity of depressive symptoms in children and adolescents. The CDRS-R consists of 17 items covering core mood, behavioral, and somatic symptoms associated with depression, each rated on a 5- or 7-point scale depending on the item. Total scores can range from 17 to 113, with higher scores indicating greater depressive symptom severity. Clinical thresholds have been established to indicate mild, moderate, and severe depression, with a score of ≥40 typically used to suggest the presence of clinically relevant depressive symptomatology.^96^ The CDRS-R has demonstrated strong psychometric properties, including high internal consistency, inter-rater reliability, and validity in both community and clinical samples of children and adolescents ^97^. In the present study, total scores were used as a dimensional measure of depressive symptom severity.

### Interoceptive accuracy

In the heartbeat counting task,^84^ participants were asked to silently count the heartbeats they perceived in their body from the moment a “start” signal was displayed on a computer screen, until a “stop” signal was displayed, without physically checking their pulse. The task consisted of three trials with time intervals of 25, 35, and 40 seconds, presented in a randomized sequence. The tasks was realized in PsychoPy (version: 1.84) ^98^ For each trial, the number of perceived heartbeats was compared to the actual number of heartbeats recorded via ECG, and an interoceptive accuracy (IAcc) score was calculated using the formula:^99^

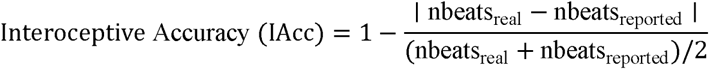

These trial scores were then averaged across the three trials to obtain a single performance score per participant. Including the reported heartbeat count in the denominator helps prevent inflated accuracy estimates in individuals with high variability or in cases where participants overestimated their heartbeats.^99^

### Physiological Recordings

The processing and analysis of cardiac vagal activity indexed by vmHRV followed the Guidelines for Reporting Articles on Psychiatry and Heart Rate Variability (GRAPH).^100^ Electrocardiographic (ECG) data were continuously recorded as interbeat intervals (IBIs) at 1,024 Hz using an ECGMove 3 sensor (movisens GmbH, Karlsruhe, Germany) attached to the chest at the base of the sternum with a flexible belt and two pre-moistened electrodes. The ECG signal was acquired during a 5-min standardized baseline period while participants performed a minimally demanding Color Detection Task (CDT),^101^ and during the heartbeat counting task. During the CDT, a colored rectangle (10 × 12 cm) changed to one of six colors (yellow, white, red, blue, green, purple) every 10 s in randomized order, and participants were instructed to count the number of occurrences of a target color. Task onset and offset were automatically logged to allow precise synchronization with ECG recordings. To ensure sufficient signal quality for HRV analysis, ECG data were visually inspected immediately after acquisition using UnisensViewer (http://unisens.org). The signals were exported in CSV format and processed in Kubios HRV 3.0 Premium.^89^ R-peaks were manually corrected, and artifacts were identified and removed. A smoothing priors detrending method (λ = 500) was applied to the IBI series, and the resulting time-domain parameters were exported in TXT format for further analysis in R using the RHRV package.^102^ From the 5-min baseline segments (CDT), the root mean square of successive differences (RMSSD) of IBIs (in milliseconds) was extracted as the primary index of vmHRV, reflecting vagally mediated cardiac control.^103^ Mean recording duration for the baseline segment was 318.93 s (SD = 1.13), and on average 1.0% of IBIs (SD = 1.46) were removed during artifact correction.

### Statistical Analysis

All statistical analyses were conducted in R Studio (v4.3.0; R Core Team).^104^ Potential sex differences in emotion regulation, depressive symptom severity, personality functioning, interoceptive accuracy (IAcc), and vagally mediated heart rate variability (vmHRV) were first examined using separate linear regression models with sex as the predictor while adjusting for age and medication status (medication vs. no medication). These analyses were conducted using the lme4 package.^90^

To examine whether difficulties in emotion regulation mediated the associations of vmHRV and IAcc with depressive symptom severity and personality functioning, structural equation modeling (SEM) was performed using the lavaan package.^91^ A multi-group SEM was estimated with biological sex specified as the grouping variable. Difficulties in emotion regulation, indexed by the Difficulties in Emotion Regulation Scale (DERS) total score, were specified as the mediator. Depressive symptom severity (CDRS-R total score) and personality (dys-)functioning (STiP-5.1 total score) were modeled simultaneously as correlated outcomes. Resting vmHRV (rMSSD), heartbeat detection accuracy (IAcc), and their interaction term (rMSSD × IAcc) were included as exogenous predictors to test both their independent and synergistic associations with emotion regulation and clinical outcomes. Medication status and age were included as covariates in the mediator model. Covariances were estimated among the exogenous predictors, and residual covariance between the two outcome variables was freely estimated.

To evaluate potential sex-specific effects, paths from rMSSD, IAcc, and their interaction to emotion regulation, as well as the corresponding direct effects on depressive symptoms and personality functioning, were estimated separately for males and females. In contrast, the associations between emotion regulation difficulties and both outcome variables were constrained to be equal across sexes. Sex-specific indirect and total effects were estimated within the SEM, and differences in indirect effects between males and females were tested directly using model-defined parameters.

Model fit was evaluated using the χ² statistic, Comparative Fit Index (CFI), Root Mean Square Error of Approximation (RMSEA), and Standardized Root Mean Square Residual (SRMR).

As a complementary analysis, each structural pathway was additionally examined using separate linear regression models including sex-by-predictor interaction terms to facilitate interpretation of sex-specific effects outside the SEM framework. Complete results of these analyses are provided in Supplementary Materials 1–3.

## 3. Results

### Descriptive Results

The final sample included in the statistical analyses comprised *N* = 279 patients (n = 211 female) with a mean age of 15.27 years (*SD* = 1.42; range: 11–18). Of these, 44.4% (*N* = 60) had graduated from school at the assessment time point. Patients had a mean BMI of 21.73 kg/m2 (*SD* = 5.11) and 74.2% (*N* = 207) of patients were taking at least one or more medications at the time of assessment. A comprehensive overview of the overall sociodemographic and clinical characteristics is provided in *Table 1*.

**Table 1.** Sample Descriptives by Sex.

| Characteristic | Sex |  |  | p-value<sup>2</sup> |
| --- | --- | --- | --- | --- |
|  | male<br>N = 67<sup>1</sup> | female<br>N = 212 | Overall<br>N = 279<sup>1</sup> |  |
| Age (years) | 15.34 (1.51) | 15.23 (1.38) | 15.28 (1.42) | 0.65 |
| BMI | 24.29 (7.12) | 21.04 (5.64) | 22.54 (5.81) | 0.33 |
| completed school<br>education |  |  |  | 0.59 |
| No | 20 / 67 (30%) | 115 / 212 (54%) | 148 / 279 (53%) |  |
| Yes | 34 / 67 (70%) | 98 / 212 (46%) | 132 / 279 (47%) |  |
| Medication use |  |  |  | 0.52 |
| No | 20 / 67 (30%) | 53 / 212 (25%) | 41 / 135 (30%) |  |
| Yes | 47 / 67 (70%) | 160 / 212 (75%) | 94 / 130 (70%) |  |
| DERS total score | 38.51 (16.88) | 52.31 (14.94) | 49.01 (16.49) | <0.001 |
| CDRS total score | 43.55 (15.84) | 54.67 (16.37) | 52.01 (16.90) | <0.001 |
| STiP total score | 12.16 (9.17) | 14.99 (8.96) | 14.31 (9.08) | 0.026 |
| IAcc score | 60.49 (31.79) | 38.23 (49.65) | 43.56 (46.94) | <0.001 |
| Resting RMSSD (ms) | 60.10 (39.78) | 55.16 (37.96) | 56.34 (38.39) | 0.36 |

<sup>1</sup>Mean (SD); n / N (%)
<sup>2</sup>One-way ANOVA; Pearson's Chi-squared test

### Sex effects

Females showed significantly more difficulties in emotion regulation (i.e. higher DERS score) compared to males, *β* = 1.04, *p* < .001. In addition, older age was associated with higher emotion dysregulation, *β* =0.14, *p* < .001, whereas medication status did not exert a significant effect. *β* = 0.17, *p* = .19. For depressive symptom severity, females also showed significantly higher scores than males, *β* =0.79, *p* < .001, and age was positively associated with symptom severity, *β* =0.15, *p* < .001, whereas medication status was not significant, *β* =0.14, *p* = .29. Similarly, personality functioning differed significantly by sex, *β* =0.36, *p* < .05, but was not significantly predicted by age, *β* = 0.04, *p* = .26 or medication status, *β* = 0.13, *p* = .34. For IAcc, females exhibited significantly lower accuracy compared to males, *β* = –0.44, *p* < .001. Neither age, *β* = –0.06, *p* = .08, nor medication status, *β* = 0.17, *p* = .18, significantly predicted IAcc. In contrast, for vmHRV, medication intake was significantly associated with reduced vmHRV, *β* = –0.55, *p* < .001, whereas no significant effects emerged for sex, *β* = -0.01, *p* = .94) or age, *β* = –0.03, *p* = .46. Sex-specific means for all variables are reported in *Table 1*, and the distributions for males and females are illustrated in *Figure 1*.

**Figure 1.**
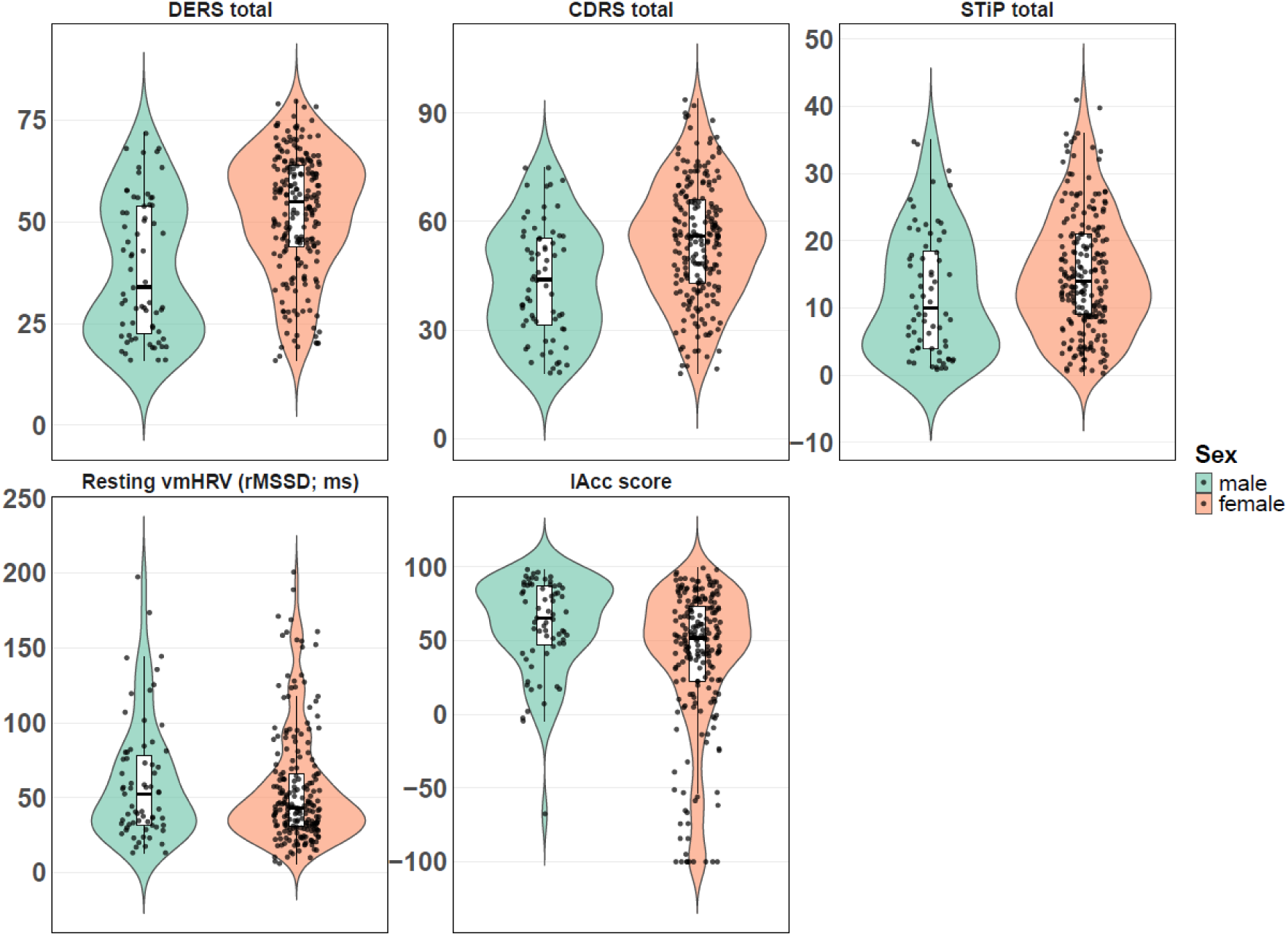
Violin plots depicting the distributions for DERS scores, CDRS scores, STiP scores (top row) and resting vmHRV, and IAcc scores (bottom row) for female and male participants, with overlaid data points and boxplots indicating median and interquartile range.

### Mediation Analyses

The mediation model showed good overall fit (χ²(6) = 9.02, *CFI* = .994, *RMSEA* = .060, *SRMR* = .017). Across sexes, higher emotion regulation difficulties were significantly associated with greater depressive symptom severity, *β* = 0.71, *p* < .001, and greater personality dysfunction, *β* = 0.31, *p* < .001.

In males, the mediation model explained 14.8%, 62.5%, and 46.2% of the variance in emotion regulation, depressive symptom severity, and personality dysfunction, respectively. Emotion dysregulation was significantly predicted by vmHRV, *β* = –0.667, *p* < .01, but not by Iacc, *β* = 0.11, *p* = .34. A significant interaction effect between Iacc and vmHRV, *β* = 0.66, *p* < .01, on the total DERS score further indicated that their combined effect was particularly relevant for emotion regulation capacity. No direct effects of Iacc and vmHRV on depressive symptoms or personality dysfunction were found (all *p*s > .05). Mediation analyses revealed significant indirect effects of vmHRV on depressive symptom severity*, β* = –0.11, *p* < .01, and personality dysfunction, *β* = –0.39, *p* < .01, through emotion dysregulation. The interaction term between vmHRV and Iacc also exerted significant indirect effects on both depressive symptom severity, *β* = 0.49, *p* <.01, and personality dysfunction, *β* = 0.38, *p* < .01. No indirect effects of Iacc on neither depressive symptoms nor personality dysfunction were found. Total effects of vmHRV, *β* = –0.65, *p* < .001, and the vmHRV x Iacc interaction, *β* = 0.54, *p* < .05, on depressive symptom severity, as well as of vmHRV, *β* = –0.77, *p* < .01, and the vmHRV x Iacc interaction, *β* = 0.57, *p* < .05, on personality dysfunction, remained significant, indicating partial mediation via emotion dysregulation. The residual covariance between depressive symptom severity and personality dysfunction remained significant after controlling for emotion regulation, *β* = 0.42, *p* < .01, indicating that emotion dysregulation did not fully account for their shared variance (see *Figure 2*). The covariance between vmHRV and Iacc was also found to be significant, *β* = 0.29, *p* < .05.

**Figure 2.**
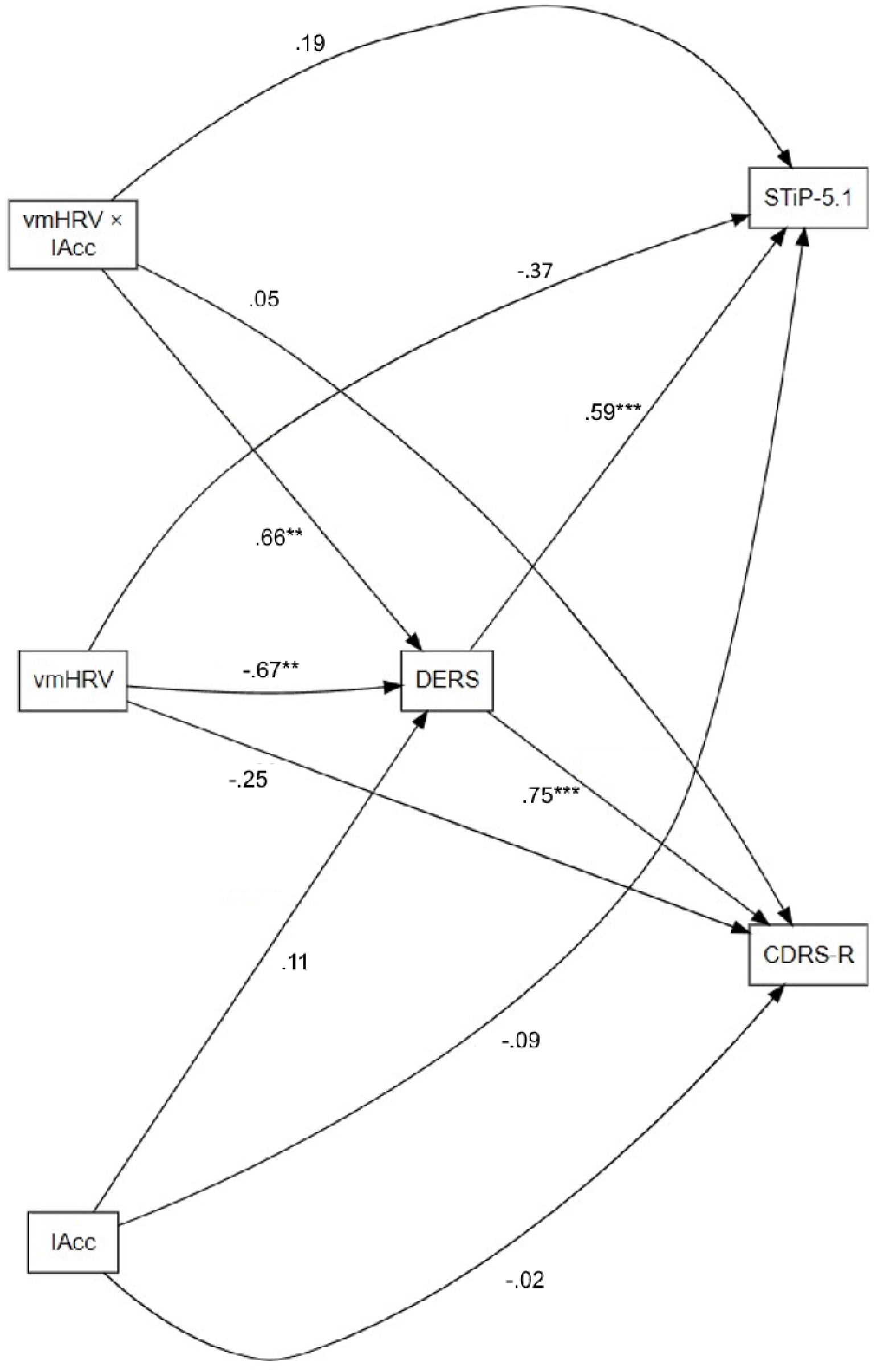
Path model depicting the mediation of interoceptive accuracy (IAcc) and vagally mediated heart rate variability (vmHRV) on depressive symptom severity (CDRS-R) and personality dysfunction (STiP-5.1) via emotion dysregulation (DERS) in the male sample. Direct paths from IAcc, vmHRV, and their interaction (vmHRV × IAcc) to DERS are shown, as well as direct paths to CDRS-R and STiP-5.1. Paths from DERS to CDRS-R and STiP-5.1 illustrate the mediating role of emotion dysregulation. Standardized regression coefficients are displayed for each path. *p < .05, **p < .01, ***p < .001. Covariates omitted for clarity.

In females, the mediation model explained 7.0%, 44.3%, and 29.0% of the variance in emotion dysregulation, depressive symptom severity, and personality dysfunction, respectively. We found a significant total effect of Iacc on personality dysfunction, *β* = 0.15, *p* < .05. No other significant direct, indirect or total effects emerged. The covariance between depressive symptom severity and personality dysfunction was significant, *β* = 0.44, *p* < .01, indicating shared variance between the outcomes not accounted for by emotion dysregulation (see *Supplementary Figure 1*). The covariance between vmHRV and Iacc was also found to be significant, *β* = 0.19, *p* < .01.

## 4. Discussion

In this study, we investigated whether cardiac vagal activity and interoceptive capacity are associated with individual differences in emotion dysregulation, a central transdiagnostic feature of psychopathology, and whether these associations extend to depressive symptoms and personality dysfunction in a large sample of adolescent psychiatric inpatients. Given well-documented sex differences in mental health outcomes, we additionally tested sex-specific effects within our proposed models.

Consistent with extensive evidence, suggesting emotion dysregulation as a transdiagnostic hallmark of psychopathology, ^3,55,56^ our analyses identified emotion dysregulation as a robust correlate of both depressive symptom severity and personality dysfunction across sexes. In addition, our analyses examining the psychophysiological correlates of emotion dysregulation revealed pronounced sex-specific patterns: in male adolescents, lower vmHRV was indirectly associated with depressive symptoms and personality dysfunction through emotion dysregulation, whereas comparable associations were absent in female adolescents. Notably, these associations were observed primarily at lower to intermediate levels of interoceptive accuracy, indicating that the relationship between cardiac vagal regulation and affective dysfunction may depend on the interplay between autonomic regulation and interoceptive processing. These findings converge with extensive empirical and theoretical accounts highlighting close links among interoceptive processing, autonomic regulation, and emotion regulation capacities that may shape vulnerability and resilience across psychiatric disorders.

Contemporary models of emotion regulation increasingly conceptualize adaptive emotional functioning as emerging from dynamic interactions between bodily and central processes. Within these frameworks, interoceptive and autonomic signals are thought to contribute to multiple stages of emotional regulation, including the perception and differentiation of emotional states, the evaluation of emotional responses, and the flexible implementation and monitoring of adaptive regulatory strategies. In line with these accounts, our findings indicate that cardiac vagal activity and interoceptive processing are jointly associated with individual differences in emotion dysregulation. Specifically, greater vmHRV was associated with fewer difficulties in emotion regulation and lower depressive symptom severity as well as personality dysfunction, particularly in males with lower interoceptive accuracy. These results suggest that higher levels of vmHRV may buffer adverse effects of low interoceptive accuracy on emotion regulation capacities. In line with this interpretation, vmHRV has previously been shown to act as a protective factor, buffering the adverse effects of early adversity^57–60^ and emotion regulation difficulties^61^ on mental health, as well as the impact of daily stress on negative affect^62^ and interpersonal stress on inflammatory responses.^63^ Similarly, higher interoceptive accuracy was linked to less emotion regulation difficulties in males with low vmHRV. Interpreting low resting vmHRV as a state of chronically elevated autonomic arousal, these findings may indicate that greater interoceptive capacity may enable more effective perception, integration, and regulation of emotions in states of heightened arousal, whereas lower interoceptive sensitivity may limit adaptive regulation. In males with higher vmHRV, however, greater interoceptive accuracy was associated with increased difficulties in emotion regulation and higher depressive symptom severity. Notably, increased weighting of interoceptive afferent input has been proposed as a pathophysiological mechanism contributing to mood and anxiety disorders.^35^ However, whereas these disorders are typically characterized by blunted vmHRV, the present associations between interoceptive accuracy, emotion dysregulation, and depressive symptom severity emerged only among individuals with comparatively high vmHRV. This apparent discrepancy may be attributable to characteristics of our clinical sample, as the psychiatric inpatient population studied here may have exhibited overall reduced vmHRV relative to healthy individuals. Nonetheless, our results reveal an interactive interplay between cardiac vagal and interoceptive processes in the (dys-) regulation of emotion and the emergence of psychopathology, emphasizing the need for integrative models and further empirical investigation.

Importantly, in our sample these interactive mechanisms emerged exclusively in male adolescents, with no comparable associations observed in females. Meta-analytic evidence has demonstrated robust sex differences in both domains, with adult women typically exhibiting higher resting cardiac vagal activity but lower interoceptive accuracy compared to men. Beyond these general distinctions, accumulating evidence further indicates that the associations between these physiological processes and mental health outcomes are themselves sex dependent. For cardiac vagal activity, sex-specific associations have been reported for emotion regulation, substance use^64^, positive^65^ and negative^66^ affect and depressive symptoms^65,67,68^. Similarly, sex differences have been observed in the relationships between interoceptive accuracy and emotion recognition^69^, emotion regulation^70–73^ and emotion perception^74–76^. Nonetheless, a recent meta-analysis synthesizing this literature found no consistent sex differences in the relationship between interoception and emotion-related outcomes and, notably, no reliable overall association between interoception and emotional measures underscoring the considerable heterogeneity in existing findings. Moreover, across both interoception and cardiac vagal activity, the directionality of reported sex differences has been inconsistent, with some studies identifying significant associations only in males and others exclusively in females.

A potential pathway underlying these sex differences involves the influence of gonadal hormones on autonomic and interoceptive processes across both short- and long-term timescales. Fluctuations in ovarian hormones such as estradiol and progesterone can modify cardiac function through peripheral hormone receptors and can also shape central autonomic regulation through their actions on regions that contribute to vagal control, including the medial prefrontal cortex, the insula, and several brainstem nuclei. Following menarche, cyclical variations in hormonal levels have been shown to not only lead to shifting levels of cardiac vagal activity but also to changing patterns of communication between the brain and the body.^77^ Such rhythmic variations may produce cycle dependent links between vagal activity, emotion regulation, and mental health, which in turn may help to explain why studies that do not account for cycle phase often report inconsistent or null findings. Similarly, monthly hormonal fluctuations have been proposed to modulate interoceptive processing. Although direct evidence is still emerging, hormonal fluctuations are believed to influence how precisely internal bodily signals are perceived and integrated.^39,78^ Supporting this interpretation, sex differences in interoceptive accuracy are typically absent before puberty^79,80^ which may also account for the absence of significant sex differences in Iacc in our sample of adolescents aged 11 to 18 years. More specifically, such hormonally driven fluctuations are hypothesized to reduce the reliability and stability of interoceptive signals as informative cues for emotional states, thereby increasing the relative influence of exteroceptive, context-related information in the generation and interpretation of emotional experiences.^81^ Our results support these accounts, indicating no significant link between interoceptive accuracy and emotion (dys-) regulation in female patients. This sex-specific dissociation raises important questions regarding the potential mechanisms that may differentially shape the integration of interoceptive and autonomic processes across sexes, thereby challenging generalized models of emotion (dys-) regulation and psychopathology that link interoception, cardiac vagal activity, and emotion regulation without considering sex-specific pathways.^11,16,82,83^ From a clinical perspective, these findings further suggest that sex-dependent differences in body–brain interactions may have implications for the development of tailored intervention approaches. In particular, they raise the possibility that the relevance of interoceptive and autonomic processes for emotion dysregulation and treatment response may vary across sexes, highlighting the need to consider sex-specific pathways when designing and targeting interventions that engage bodily signals in the regulation of affect.

Taken together, our findings underscore the importance of jointly considering interoceptive and autonomic pathways when characterizing the mechanisms that shape emotion (dys-)regulation and confer vulnerability to psychopathology. Crucially, the marked sex-specificity of these associations highlights that models developed primarily from mixed or predominantly adult samples may obscure distinct developmental trajectories in males and females. By demonstrating interactive contributions of cardiac vagal and interoceptive processes only in male adolescents, our results call for theoretical and clinical frameworks that explicitly incorporate sex as a biological and developmental moderator rather than treating it as a nuisance variable. Future work integrating longitudinal, hormonal, and neurobiological measures will be essential for delineating how these systems co-develop and how their interactions give rise to sex-differentiated pathways of risk and resilience.

Several limitations should be considered when interpreting these findings. First, interoception was assessed exclusively using the heartbeat counting task,^84^ a measure that has been criticized for its susceptibility to prior knowledge about one’s heart rate and for capturing only a single facet of interoception, i.e., interoceptive accuracy, while omitting related dimensions such as interoceptive sensibility and interoceptive awareness. ^85,86^ Second, although hormonal fluctuations are central to many proposed mechanisms underlying sex differences in interoceptive and autonomic functioning, our study did not include information on menarcheal status, menstrual cycle phase, or other indicators of hormonal milieu, limiting our ability to directly evaluate hormone-related effects. Third, because the sample consisted solely of psychiatric inpatients, the generalizability of the observed associations to community or non-clinical populations remains uncertain. Fourth, the findings regarding the moderating effect of sex are limited by unequal subsample sizes (*n* = 212 females; *n* = 67 males). However, unequal sex distributions are common in clinical samples, and the present sample reflects this common pattern. Lastly, although our moderated mediation model was theoretically grounded in contemporary models of body–brain interactions and emotion (dys-) regulation, the cross-sectional design precludes conclusions regarding causal directionality or temporal ordering among variables. While the proposed model is consistent with the assumption that cardiac vagal activity and interoceptive processing influence emotion (dys-)regulation, which in turn relates to psychopathological outcomes, alternative or reciprocal pathways are equally plausible. Future longitudinal and experimental studies are therefore required to clarify the temporal sequencing and mechanistic pathways underlying these associations. Furthermore, future research should extend these findings by employing multi-dimensional interoception measures, incorporating detailed hormonal assessments, and examining more diverse samples to determine the scope and boundaries of the reported sex-specific pathways.

## Resource availability

### Lead contact

Further information and requests for resources should be directed to and will be fulfilled by the lead contact, Julian Koenig.

### Materials availability

This study did not generate any new, unique reagents.

### Data and code availability

- The data reported in this paper will be shared by the lead contact upon request.
- All original code used for the statistical analyses reported in this study has been deposited via OSF (https://osf.io/rxutn/overview?view_only=f4b7cd804cfc4744a4d837c186c3f731)
- Any additional information required to reanalyze the data reported in this paper is available from the lead contact upon request.

## Acknowledgements

Julian Koenig acknowledges financial support for the Mapping Autonomic Neural Interaction and Control (MANIAC) Emerging Group by the University of Cologne Excellent Research Support Program.

## Author contributions

M.K. and J.K. conceptualized the study; H.S., I.M.-L., C.R., M.K., and J.K. acquired data; M.S. analyzed the data; M.S. and J.K. interpreted the findings; M.S. drafted the manuscript; and H.S., I.M.-L., C.R., M.K., and J.K. reviewed and edited the manuscript. All authors approved the final version of the manuscript.

## Declaration of Interests

The authors have no conflict of interest to report.

## Declaration of generative AI and AI-assisted technologies in the writing process

During the preparation of this work, the authors used ChatGPT-4o in order to improve clarity, refine language, and check grammar. After using this tool or service, the authors reviewed and edited the content as needed and take full responsibility for the content of the publication.

## Supplementary Material

### Supplementary Material 1 - Effects of Interoceptive Accuracy and vagally mediated HRV on Emotion dysregulation

A multiple regression model was conducted to examine the combined and interacting effects of interoceptive accuracy (IAcc), vagally mediated heart rate variability (vmHRV), and sex on emotion dysregulation (DERS), controlling for age and medication status. A significant main effect emerged for vmHRV, *β* = –0.61, *p* < .01, and sex (female), *β* = 1.03, *p* < .001. Significant two-way interactions were observed for IAcc × vmHRV, *β* = 0.77, *p* < .01 and vmHRV × sex, *β* = 0.71, *p* < .01. The three-way interaction of IAcc × vmHRV × sex was also significant, *β* = –0.78, *p* < .01. The full model explained 18.8% of the variance in DERS scores, *F*(9,269) = 8.13, *p* < .001.

Post-hoc simple slopes analyses in males indicated that higher vmHRV predicted lower DERS scores at low, *β* = –1.31, *p* < .001, and mean, *β* = –0.58, *p* < .05, levels of IAcc, but not at high IAcc, *β* = 0.16, *p* = .24. Similarly, IAcc was associated with DERS scores at high vmHRV, *β* = 0.81, *p* < .05, with no effect at low, *β* = -0.61, *p* = .06, and mean vmHRV, *β* = 0.20, *p* = 0.27. In females, none of the simple slopes reached significance.

### Supplementary Material 2 - Effects of Interoceptive Accuracy and vagally mediated HRV on Personality Functioning (STiP-5.1)

A multiple regression model was conducted to examine the combined and interacting effects of IAcc, vmHRV, and sex on personality dysfunction (STiP-5.1), controlling for age and medication status. A significant main effect emerged for vmHRV, *β* = –0.69, *p* < .01. While there were significant two-way interactions between IAcc and vmHRV, *β* = 0.66, *p* < .05, and between vmHRV and sex, *β* = 0.65, *p* < .01, the three-way interaction of IAcc × vmHRV × sex, *β* = –0.57, *p* = .05, did not reach significance. The full model explained 4.9 % of the variance in STiP-5.1 scores, *F*(9,269) = 2.58, *p* = .007.

Post-hoc simple slopes analyses in males indicated that higher vmHRV predicted lower STiP-5.1 scores at low, *β* = –1.29, *p* < .05, and mean, *β* = –0.66, *p* < .01, levels of IAcc, but not at high IAcc, *p* = .86. Similarly, IAcc was associated with STiP-5.1 scores at low vmHRV, *β* = –0.73, *p* < .05, with no significant effects at mean or high vmHRV. In females, none of the simple slopes reached significance.

### Supplementary Material 3 - Effects of Interoceptive Accuracy and vagally mediated HRV on Depressive Symptom Severity (CDRS-R)

A multiple regression model was conducted to examine the combined and interacting effects of IAcc, vmHRV, and sex on depressive symptom severity (CDRS-R), controlling for age and medication status. Significant main effects emerged for vmHRV, *β* = –0.62, *p* < .01, and sex (female), *β* = 0.78, *p* < .001. Significant two-way interactions were observed for IAcc × vmHRV, *β* = 0.58, *p* < .05, and vmHRV × sex, *β* = 0.63, *p* < .01. The three-way interaction of IAcc × vmHRV × sex was also significant, *β* = –0.57, *p* < .01. The full model explained 14.3% of the variance in CDRS-R scores, *F*(9,269) = 6.13, *p* < .001.

Post-hoc simple slopes analyses in males indicated that higher vmHRV predicted lower CDRS-R scores at low, *β* = –1.15, *p* < .05, and mean, *β* = –0.60, *p* < .01, levels of IAcc, but not at high IAcc, *p* = .74. Similarly, IAcc was associated with CDRS-R scores at high vmHRV, *β* = 0.68, *p* < .05, with no effect at low and mean vmHRV. In females, none of the simple slopes were significant.

---

**Supplementary Figure 1.**
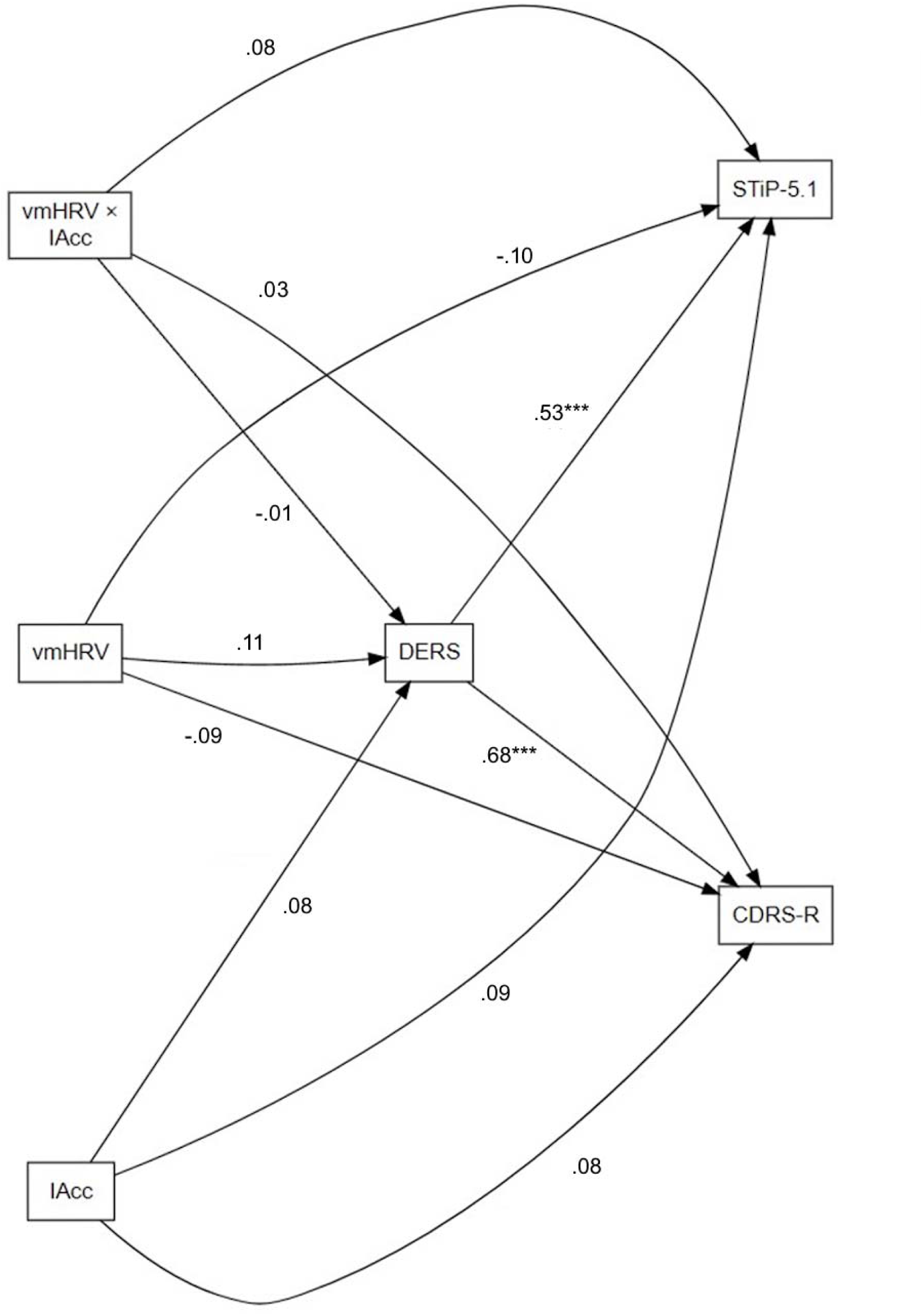
Path model depicting the mediation of interoceptive accuracy (IAcc) and vagally mediated heart rate variability (vmHRV) on depressive symptom severity (CDRS-R) and personality dysfunction (STiP-5.1) via emotion dysregulation (DERS) in the female sample. Direct paths from IAcc, vmHRV, and their interaction (vmHRV × IAcc) to DERS are shown, as well as direct paths to CDRS-R and STiP-5.1. Paths from DERS to CDRS-R and STiP-5.1 illustrate the mediating role of emotion dysregulation. Standardized regression coefficients are displayed for each path. *p < .05, **p < .01, ***p < .001.

## Notes

### Competing Interest Statement

The authors have declared no competing interest.

## References

1. Beauchaine, T. P. & Cicchetti, D. Emotion dysregulation and emerging psychopathology: A transdiagnostic, transdisciplinary perspective. Development and Psychopathology 31, 799–804 (2019).

2. Beauchaine, T. P. Future Directions in Emotion Dysregulation and Youth Psychopathology. Journal of Clinical Child & Adolescent Psychology 44, 875–896 (2015).

3. Sheppes, G., Suri, G. & Gross, J. J. Emotion Regulation and Psychopathology. Annual Review of Clinical Psychology 11, 379–405 (2015).

4. Gratz, K. L. & Roemer, L. Multidimensional Assessment of Emotion Regulation and Dysregulation: Development, Factor Structure, and Initial Validation of the Difficulties in Emotion Regulation Scale. Journal of Psychopathology and Behavioral Assessment 26, 41–54 (2004).

5. Faustino, B. Transdiagnostic perspective on psychological inflexibility and emotional dysregulation. Behavioural and Cognitive Psychotherapy 49, 233–246 (2021).

6. Chin, M., Robson, D. A., Woodbridge, H. & Hawes, D. J. Irritability as a Transdiagnostic Construct Across Childhood and Adolescence: A Systematic Review and Meta-analysis. Clin Child Fam Psychol Rev 28, 101–124 (2025).

7. Wolff, J. C. et al. Emotion dysregulation and non-suicidal self-injury: A systematic review and meta-analysis. European Psychiatry 59, 25–36 (2019).

8. Hallion, L. S., Steinman, S. A., Tolin, D. F. & Diefenbach, G. J. Psychometric Properties of the Difficulties in Emotion Regulation Scale (DERS) and Its Short Forms in Adults With Emotional Disorders. Front. Psychol. 9, (2018).

9. Thayer, J. F. & Lane, R. D. A model of neurovisceral integration in emotion regulation and dysregulation. J Affect Disord 61, 201–216 (2000).

10. Barrett, L. F., Quigley, K. S. & Hamilton, P. An active inference theory of allostasis and interoception in depression. Philos Trans R Soc Lond B Biol Sci 371, 20160011 (2016).

11. Seth, A. K. & Critchley, H. D. Extending predictive processing to the body: Emotion as interoceptive inference. Behav Brain Sci 36, 227–228 (2013).

12. Craig, A. D. How do you feel? Interoception: the sense of the physiological condition of the body. Nat Rev Neurosci 3, 655–666 (2002).

13. Ainley, V., Apps, M. A. J., Fotopoulou, A. & Tsakiris, M. ‘Bodily precision’: a predictive coding account of individual differences in interoceptive accuracy. Philosophical Transactions of the Royal Society B: Biological Sciences 371, 20160003 (2016).

14. Khalsa, S. S., Rudrauf, D., Feinstein, J. S. & Tranel, D. The pathways of interoceptive awareness. Nat Neurosci 12, 1494–1496 (2009).

15. Owens, A. P., Allen, M., Ondobaka, S. & Friston, K. J. Interoceptive inference: From computational neuroscience to clinic. Neuroscience & Biobehavioral Reviews 90, 174–183 (2018).

16. Critchley, H. D. & Garfinkel, S. N. Interoception and emotion. Current Opinion in Psychology 17, 7–14 (2017).

17. Feldman, M. J., Bliss-Moreau, E. & Lindquist, K. A. The neurobiology of interoception and affect. Trends in Cognitive Sciences 28, 643–661 (2024).

18. Greenwood, B. M. & Garfinkel, S. N. Interoceptive Mechanisms and Emotional Processing. Annual Review of Psychology 76, 59–86 (2025).

19. Zamariola, G., Frost, N., Van Oost, A., Corneille, O. & Luminet, O. Relationship between interoception and emotion regulation: New evidence from mixed methods. J Affect Disord 246, 480–485 (2019).

20. Zsembik, L., Oldroyd, P. & Chen, R. Interoceptive modulation of emotions. Current Opinion in Neurobiology 92, 103049 (2025).

21. Barrett, L. F., Quigley, K. S., Bliss-Moreau, E. & Aronson, K. R. Interoceptive Sensitivity and Self-Reports of Emotional Experience. Journal of Personality and Social Psychology 87, 684–697 (2004).

22. Dunn, B. D. et al. Listening to Your Heart: How Interoception Shapes Emotion Experience and Intuitive Decision Making. Psychol Sci 21, 1835–1844 (2010).

23. Ventura-Bort, C., Wendt, J. & Weymar, M. The Role of Interoceptive Sensibility and Emotional Conceptualization for the Experience of Emotions. Front. Psychol. 12, (2021).

24. Füstös, J., Gramann, K., Herbert, B. M. & Pollatos, O. On the embodiment of emotion regulation: interoceptive awareness facilitates reappraisal. Soc Cogn Affect Neurosci 8, 911–917 (2013).

25. Gray, M. A. et al. Emotional appraisal is influenced by cardiac afferent information. Emotion 12, 180–191 (2012).

26. Kever, A., Pollatos, O., Vermeulen, N. & Grynberg, D. Interoceptive sensitivity facilitates both antecedent- and response-focused emotion regulation strategies. Personality and Individual Differences 87, 20–23 (2015).

27. Arai, T., Komano, T., Munakata, T. & Ohira, H. The association between interoception and olfactory affective responses. Biological Psychology 193, 108878 (2024).

28. Handy, A. B., Freihart, B. K. & Meston, C. M. The Relationship between Subjective and Physiological Sexual Arousal in Women with and without Arousal Concerns. Journal of Sex & Marital Therapy 46, 447–459 (2020).

29. MacCormack, J. K., Bonar, A. S. & Lindquist, K. A. Interoceptive beliefs moderate the link between physiological and emotional arousal during an acute stressor. Emotion 24, 269–290 (2024).

30. Brewer, R., Cook, R. & Bird, G. Alexithymia: a general deficit of interoception. Royal Society Open Science 3, 150664 (2016).

31. Herbert, B. M., Herbert, C. & Pollatos, O. On the Relationship Between Interoceptive Awareness and Alexithymia: Is Interoceptive Awareness Related to Emotional Awareness? Journal of Personality 79, 1149–1175 (2011).

32. Khalsa, S. S. et al. Interoception and Mental Health: A Roadmap. Biol Psychiatry Cogn Neurosci Neuroimaging 3, 501–513 (2018).

33. Lavalley, C. A. et al. Transdiagnostic failure to adapt interoceptive precision estimates across affective, substance use, and eating disorders: A replication study. medRxiv 2023.10.11.23296870 (2023) doi:10.1101/2023.10.11.23296870.

34. Nord, C. L. & Garfinkel, S. N. Interoceptive pathways to understand and treat mental health conditions. Trends in Cognitive Sciences 26, 499–513 (2022).

35. Paulus, M. P., Feinstein, J. S. & Khalsa, S. S. An Active Inference Approach to Interoceptive Psychopathology. Annual Review of Clinical Psychology 15, 97–122 (2019).

36. Paulus, M. P. & Stein, M. B. Interoception in anxiety and depression. Brain Struct Funct 214, 451–463 (2010).

37. Smith, A., Forrest, L. & Velkoff, E. Out of touch: Interoceptive deficits are elevated in suicide attempters with eating disorders. Eating Disorders 26, 52–65 (2018).

38. Back, S. N. & Bertsch, K. Interoceptive Processing in Borderline Personality Pathology: a Review on Neurophysiological Mechanisms. Curr Behav Neurosci Rep 7, 232–238 (2020).

39. Murphy, J., Viding, E. & Bird, G. Does atypical interoception following physical change contribute to sex differences in mental illness? Psychol Rev 126, 787–789 (2019).

40. Appelhans, B. M. & Luecken, L. J. Heart Rate Variability as an Index of Regulated Emotional Responding. Review of General Psychology 10, 229–240 (2006).

41. Kreibig, S. D. Autonomic nervous system activity in emotion: A review. Biological Psychology 84, 394–421 (2010).

42. Wu, Y., Gu, R., Yang, Q. & Luo, Y. How Do Amusement, Anger and Fear Influence Heart Rate and Heart Rate Variability? Front. Neurosci. 13, (2019).

43. Hoemann, K. et al. Investigating the relationship between emotional granularity and cardiorespiratory physiological activity in daily life. Psychophysiology 58, e13818 (2021).

44. Grol, M. & De Raedt, R. The link between resting heart rate variability and affective flexibility. Cogn Affect Behav Neurosci 20, 746–756 (2020).

45. Svendsen, J. L. et al. Trait Self-Compassion Reflects Emotional Flexibility Through an Association with High Vagally Mediated Heart Rate Variability. Mindfulness 7, 1103–1113 (2016).

46. Berna, G., Ott, L. & Nandrino, J.-L. Effects of Emotion Regulation Difficulties on the Tonic and Phasic Cardiac Autonomic Response. PLOS ONE 9, e102971 (2014).

47. Visted, E. et al. The Association between Self-Reported Difficulties in Emotion Regulation and Heart Rate Variability: The Salient Role of Not Accepting Negative Emotions. Front. Psychol. 8, (2017).

48. Williams, D. P. et al. Resting heart rate variability predicts self-reported difficulties in emotion regulation: a focus on different facets of emotion regulation. Front. Psychol. 6, (2015).

49. Cheng, Y.-C., Su, M.-I., Liu, C.-W., Huang, Y.-C. & Huang, W.-L. Heart rate variability in patients with anxiety disorders: A systematic review and meta-analysis. Psychiatry and Clinical Neurosciences 76, 292–302 (2022).

50. Kemp, A. H. et al. Impact of Depression and Antidepressant Treatment on Heart Rate Variability: A Review and Meta-Analysis. Biological Psychiatry 67, 1067–1074 (2010).

51. Koch, C., Wilhelm, M., Salzmann, S., Rief, W. & Euteneuer, F. A meta-analysis of heart rate variability in major depression. Psychological Medicine 49, 1948–1957 (2019).

52. Koenig, J., Kemp, A. H., Feeling, N. R., Thayer, J. F. & Kaess, M. Resting state vagal tone in borderline personality disorder: A meta-analysis. Progress in Neuro-Psychopharmacology and Biological Psychiatry 64, 18–26 (2016).

53. Koenig, J. & Thayer, J. F. Sex differences in healthy human heart rate variability: A meta-analysis. Neuroscience & Biobehavioral Reviews 64, 288–310 (2016).

54. Koenig, J., Rash, J. A., Campbell, T. S., Thayer, J. F. & Kaess, M. A Meta-Analysis on Sex Differences in Resting-State Vagal Activity in Children and Adolescents. Front. Physiol. 8, (2017).

55. Lincoln, T. M., Schulze, L. & Renneberg, B. The role of emotion regulation in the characterization, development and treatment of psychopathology. Nat Rev Psychol 1, 272–286 (2022).

56. Gross, J. J. & Jazaieri, H. Emotion, Emotion Regulation, and Psychopathology: An Affective Science Perspective. Clinical Psychological Science 2, 387–401 (2014).

57. Susman, E. S., Weissman, D. G., Sheridan, M. A. & McLaughlin, K. A. High Vagal Tone and Rapid Extinction Learning as Potential Transdiagnostic Protective Factors Following Childhood Violence Exposure. Dev Psychobiol 63, e22176 (2021).

58. McLaughlin, K. A., Alves, S. & Sheridan, M. A. Vagal Regulation and Internalizing Psychopathology among Adolescents Exposed to Childhood Adversity. Dev Psychobiol 56, 1036– 1051 (2014).

59. El-Sheikh, M., Harger, J. & Whitson, S. M. Exposure to interparental conflict and children’s adjustment and physical health: the moderating role of vagal tone. Child Dev 72, 1617–1636 (2001).

60. McLaughlin, K. A., Rith-Najarian, L., Dirks, M. A. & Sheridan, M. A. Low vagal tone magnifies the association between psychosocial stress exposure and internalizing psychopathology in adolescents. J Clin Child Adolesc Psychol 44, 314–328 (2015).

61. Fantini-Hauwel, C., Batselé, E., Gois, C. & Noel, X. Emotion Regulation Difficulties Are Not Always Associated With Negative Outcomes on Women: The Buffer Effect of HRV. Front. Psychol. 11, (2020).

62. da Estrela, C., MacNeil, S. & Gouin, J.-P. Heart rate variability moderates the between- and within-person associations between daily stress and negative affect. International Journal of Psychophysiology 162, 79–85 (2021).

63. Michels, N. et al. Interpersonal stressors predicting inflammation in adolescents: Moderation by emotion regulation and heart rate variability? Biological Psychology 193, 108900 (2024).

64. Kwon, E. S. et al. Resting Heart Rate Variability, Perceived Emotion Regulation, and Low-Risk Drug Use in College-Aged Adults: Gender as a Moderator. Front Psychiatry 13, 885217 (2022).

65. Spangler, D. P. et al. Gender Matters: Nonlinear Relationships Between Heart Rate Variability and Depression and Positive Affect. Front. Neurosci. 15, (2021).

66. Verkuil, B. et al. Gender differences in the impact of daily sadness on 24-h heart rate variability. Psychophysiology 52, 1682–1688 (2015).

67. Voss, A., Boettger, M. K., Schulz, S., Gross, K. & Bär, K.-J. Gender-dependent impact of major depression on autonomic cardiovascular modulation. Progress in Neuro-Psychopharmacology and Biological Psychiatry 35, 1131–1138 (2011).

68. Thayer, J. F., Smith, M., Rossy, L. A., Sollers, J. J. & Friedman, B. H. Heart period variability and depressive symptoms: gender differences. Biological Psychiatry 44, 304–306 (1998).

69. Murphy, J., Millgate, E., Geary, H., Catmur, C. & Bird, G. No effect of age on emotion recognition after accounting for cognitive factors and depression. Q J Exp Psychol (Hove) 72, 2690–2704 (2019).

70. Lischke, A., Pahnke, R., Mau-Moeller, A., Jacksteit, R. & Weippert, M. Sex-Specific Relationships Between Interoceptive Accuracy and Emotion Regulation. Front. Behav. Neurosci. 14, (2020).

71. Jakubczyk, A. et al. Association Between Interoception and Emotion Regulation in Individuals With Alcohol Use Disorder. Front Psychiatry 10, 1028 (2019).

72. Van ’t Wout, M., Faught, S. & Menino, D. Does interoceptive awareness affect the ability to regulate unfair treatment by others? Front. Psychol. 4, (2013).

73. Schuette, S. A., Zucker, N. L. & Smoski, M. J. Do interoceptive accuracy and interoceptive sensibility predict emotion regulation? Psychol Res 85, 1894–1908 (2021).

74. Bornemann, B. & Singer, T. Taking time to feel our body: Steady increases in heartbeat perception accuracy and decreases in alexithymia over 9 months of contemplative mental training. Psychophysiology 54, 469–482 (2017).

75. Mul, C.-L., Stagg, S. D., Herbelin, B. & Aspell, J. E. The Feeling of Me Feeling for You: Interoception, Alexithymia and Empathy in Autism. J Autism Dev Disord 48, 2953–2967 (2018).

76. Soker-Elimaliah, S. et al. Autistic Traits Moderate Relations Between Cardiac Autonomic Activity, Interoceptive Accuracy, and Emotion Processing in College Students. Int J Psychophysiol 155, 118–126 (2020).

77. Prinsen, J., Villringer, A. & Sacher, J. The monthly rhythm of the brain-heart connection. Sci Adv 11, eadt1243.

78. Pennebaker, J. W. & Roberts, T.-A. Toward a His and Hers Theory of Emotion: Gender Differences in Visceral Perception. Journal of Social and Clinical Psychology 11, 199–212 (1992).

79. Schaan, L. et al. Interoceptive accuracy, emotion recognition, and emotion regulation in preschool children. International Journal of Psychophysiology 138, 47–56 (2019).

80. Koch, A. & Pollatos, O. Cardiac sensitivity in children: Sex differences and its relationship to parameters of emotional processing. Psychophysiology 51, 932–941 (2014).

81. Prentice, F., Hobson, H., Spooner, R. & Murphy, J. Gender differences in interoceptive accuracy and emotional ability: An explanation for incompatible findings. Neuroscience & Biobehavioral Reviews 141, 104808 (2022).

82. Barrett, L. F. The theory of constructed emotion: an active inference account of interoception and categorization. Soc Cogn Affect Neurosci 12, 1–23 (2017).

83. Thayer, J. F., Hansen, A. L., Saus-Rose, E. & Johnsen, B. H. Heart Rate Variability, Prefrontal Neural Function, and Cognitive Performance: The Neurovisceral Integration Perspective on Self-regulation, Adaptation, and Health. Annals of Behavioral Medicine 37, 141–153 (2009).

84. Schandry, R. Heart Beat Perception and Emotional Experience. Psychophysiology 18, 483–488 (1981).

85. Ferentzi, E., Wilhelm, O. & Köteles, F. What counts when heartbeats are counted. Trends in Cognitive Sciences 26, 832–835 (2022).

86. Garfinkel, S. N., Seth, A. K., Barrett, A. B., Suzuki, K. & Critchley, H. D. Knowing your own heart: Distinguishing interoceptive accuracy from interoceptive awareness. Biological Psychology 104, 65–74 (2015).

87. Harris, P. A. et al. The REDCap consortium: Building an international community of software platform partners. Journal of Biomedical Informatics 95, 103208 (2019).

88. Peirce, J. et al. PsychoPy2: Experiments in behavior made easy. Behav Res 51, 195–203 (2019).

89. Tarvainen, M. P., Niskanen, J.-P., Lipponen, J. A., Ranta-Aho, P. O. & Karjalainen, P. A. Kubios HRV--heart rate variability analysis software. Comput Methods Programs Biomed 113, 210–220 (2014).

90. Bates, D., et al. lme4: Linear Mixed-Effects Models using ‘Eigen’ and S4. (2026).

91. Rosseel, Y., et al. lavaan: Latent Variable Analysis. (2026).

92. Hutsebaut, J., Kamphuis, J. H., Feenstra, D. J., Weekers, L. C. & De Saeger, H. Assessing DSM-5-oriented level of personality functioning: Development and psychometric evaluation of the Semi-Structured Interview for Personality Functioning DSM-5 (STiP-5.1). Personal Disord 8, 94–101 (2017).

93. Fossati, A. & Somma, A. The assessment of personality pathology in adolescence from the perspective of the Alternative DSM-5 Model for Personality Disorder. Current Opinion in Psychology 37, 39–43 (2021).

94. Morey, L. C., Bender, D. S. & Skodol, A. E. Validating the proposed diagnostic and statistical manual of mental disorders, 5th edition, severity indicator for personality disorder. J Nerv Ment Dis 201, 729–735 (2013).

95. Zettl, M., Taubner, S., Hutsebaut, J. & Volkert, J. Psychometrische Evaluation der deutschen Version des Semistrukturierten Interviews zur Erfassung der DSM-5 Persönlichkeitsfunktionen (STiP-5.1). Psychother Psychosom Med Psychol 69, 499–504 (2019).

96. Poznanski, E. O. et al. Children’s Depression Rating Scale--Revised. 10.1037/t55280-000 (2017).

97. Mayes, T. L., Bernstein, I. H., Haley, C. L., Kennard, B. D. & Emslie, G. J. Psychometric Properties of the Children’s Depression Rating Scale–Revised in Adolescents. Journal of Child and Adolescent Psychopharmacology 20, 513–516 (2010).

98. Peirce, J. W. PsychoPy--Psychophysics software in Python. J Neurosci Methods 162, 8–13 (2007).

99. Hart, N., McGowan, J., Minati, L. & Critchley, H. D. Emotional Regulation and Bodily Sensation: Interoceptive Awareness Is Intact in Borderline Personality Disorder. Journal of Personality Disorders 27, 506–518 (2013).

100. Quintana, D. S., Alvares, G. A. & Heathers, J. a. J. Guidelines for Reporting Articles on Psychiatry and Heart rate variability (GRAPH): recommendations to advance research communication. Transl Psychiatry 6, e803 (2016).

101. Jennings, J. R., Kamarck, T., Stewart, C., Eddy, M. & Johnson, P. Alternate Cardiovascular Baseline Assessment Techniques: Vanilla or Resting Baseline. Psychophysiology 29, 742–750 (1992).

102. García Martínez, C. A., et al. Heart Rate Variability Analysis with the R Package RHRV. (Springer International Publishing, Cham, 2017). doi:10.1007/978-3-319-65355-6.

103. Malik, M. et al. Heart rate variability: Standards of measurement, physiological interpretation, and clinical use. Eur Heart J 17, 354–381 (1996).

104. R Core Team. R: A language and environment for statistical computing. R Foundation for Statistical Computing. (2022).

